# Making the Most of Potential: Potential Games and Genotypic Convergence

**DOI:** 10.1101/2020.12.08.416743

**Authors:** OMER Edhan, ZIV Hellman, ILAN Nehama

## Abstract

We consider genotypic convergence of populations and show that under fixed fitness asexual and haploid sexual populations attain monomorphic convergence (even under genetic linkage between loci) to basins of attraction with locally exponential convergence rates; the same convergence obtains in single locus diploid sexual reproduction but to polymorphic populations. Furthermore, we show that there is a unified underlying theory underlying these convergences: all of them can be interpreted as instantiations of players in a potential game implementing a multiplicative weights updating algorithm to converge to equilibrium, making use of the Baum–Eagon Theorem. To analyse varying environments, we introduce the concept of ‘virtual convergence’, under which, even if fixation is not attained, the population nevertheless achieves the fitness growth rate it would have had under convergence to an optimal genotype. Virtual convergence is attained by asexual, haploid sexual, and multi-locus diploid reproducing populations, even if environments vary arbitrarily. We also study conditions for true monomorphic convergence in asexually reproducing populations in varying environments.

## 1. Introduction

One of the central questions of evolutionary theory has long been identifying conditions for asymptotic convergence to fixation on a monomorphic population. The classical example of such a result is the simplest case of asexual reproduction without mutation (e.g., bacteria reproducing in a petri dish) in which a version of the fundamental theorem of natural selection obtains: the mean fitness of the population, which follows the dynamic of the replicator equation, increases monotonically, leading to asymptotic fixation to a monomorphic population consisting of an optimal genotype with respect to the fitness environment.

Even this strong result, however, fails to hold once one considers arbitrarily varying fitness environments over time, even in asexually reproducing populations; in sexually reproducing populations the matter is more complicated still. In this paper we consider the general question of genotypic convergence of populations implementing various reproductive strategies under conditions of both fixed and varying environments. To this end we also introduce a concept that we term ‘virtual convergence’, applying ideas originally developed for the study of algorithms.

In greater detail, we consider here three discrete time population reproductive strategies: asexual, haploid sexual, and diploid sexual. The relevant state spaces for all of these is a polytope Θ. In the asexual case Θ = Δ(Γ), the space of probability distributions over the set Γ of possible genotypes; what is of interest is tracing over time the relative frequency of the genotypes. In the sexual cases the focus instead is on the relative frequency in the population of alleles at each locus; if there are *m* loci with *k* + 1 alleles per locus, the polytope of interest is 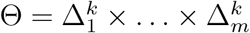

The dynamics considered will in general be describable by a transformation *T* : Θ → Θ. That is, if the population is at state *θ* ∈ Θ at time *t*, under the model it will be in state *T* (*θ*) at time *t* + 1. The main matter studied is then the asymptotics of the trajectory defined by *T^n^*(*θ*), starting from any *θ*, as *n* increases. If for each initial point *x* ∈ Θ there is a point *y* ∈ Θ such that lim_*n→∞*_ T^n^(*x*) = *y* then the dynamic converges polymorphically; if *y* is a point distribution in 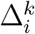 for each 1 ≤ *i* ≤ *m* then the convergence is monomorphic.

### 1.1. Fixed Environments and Convergence in Potential Games

The first question we consider asks which of these dynamics is guaranteed to converge, either monmorphically or polymorphically, when environments are fixed and unchanging over time, and we show that the asexual replicator dynamic, the sexual haploid dynamic – even under genetic inheritance linkage between loci – and the single-locus diploid dynamic all converge.

Furthermore, we provide a unified explanation for the convergence of all of these dynamics in the discrete time setting: all of them may be considered to be manifestations of potential games in which the players monotonically increase the mean potential payoff by application of a multiplicative weights updating algorithm and exploiting the Baum–Eagon Theorem. When analysing discrete time dynamics, the standard tools of continuous time gradient climbing, which depend on partial derivatives, are not available. The Baum–Eagon inequality (see Section A), which was originally intended for application to the study of hidden Markov models, provides an alternative tool that has proven to be extremely relevant to analysing discrete time evolutionary models.

The Baum–Eagon Theorem is also an essential element in our proof of a general theorem that players in a potential game independently implementing polynomial multiplicative weights updating algorithms will asymptotically converge to a fixed point that is a Nash equilibrium. The surprising element of this theorem is that monotonic climbing of the mean potential payoff is attained even though there is no coordinating element to the updating of the players, each of whom updates based on the private information of the stage payoff received without explicitly taking into account the payoffs and updated distributions of the other players.

This is especially pertinent to our study of convergence in the sexual haploid model, where the dynamic can be described as an identical interests game being played by the loci, with the objective being identifying an optimal genotype; the theorem shows that the replicator dynamic conducted independently amongst the alleles at each locus, which is the essence of the sexual reproduction model, is guaranteed to converge. The exception to all this is the multi-locus diploid model under linkage disequilibrium, where the disequilibrium term prevents application of the Baum–Eagon inequality, and in fact it has long been known that convergence under that model is not guaranteed.

A further advantage of undergirding the fixed environment theorems by appeal to the Baum–Eagon Theorem is that it enables us to make use of theorems from [Baum and Sell, 1968] to obtain finer resolution insights into the dynamic paths followed by population along the way towards convergence. This includes the fact that surrounding each pure Nash equilibrium there exists a basin of attraction, and even more strongly a basin of attraction that is exponentially stable. This implies that an observer following a path through the state space (including that of any potential game in which the players are implementing the polynomial multiplicative weights updating algorithm) will for a long time register relatively small increased in mean payoff until the path enters the exponental basin of attraction, at which point an acceleration will be noted with exponentially fast convergence to a fixed point of local maximal mean payoff.

### 1.2. Varying Environments and Virtual Convergence

When environments vary, sufficiently wildly varying environments from one time period to the next can make it impossible for the dynamic to converge to any single population state in Θ. To contend with this we introduce here a new concept of ‘virtual convergence’. This is defined using tools borrowed from computer science and introduced to the population genetics literature in the past decade, which involve regret minimisation algorithms. The metaphor often used to describe this approach is that of selecting an action with respect to varying payoff functions in subsequent time periods based on advices offered by a collection of experts. The objective is attaining asymptotically the payoff that would have been achieved had one followed from the start the advice of the best expert in hindsight in every time period; a no regret algorithm achieves this objective.

In the evolutionary setting, the analogues of the experts of the previous paragraph are genotypes and the payoffs are fitness values. The question then becomes: is it the case that, no matter what sequence of environments and hence fitness values is realised, the reproducing population asymptotically attains the mean fitness that is the growth rate that it would have achieved had it been comprised from the start monomorphically by the optimal-in-hindsight genotype? If yes, then we say that virtual convergence is attained.

With these definitions, we study here modes of convergence for the asexual, haploid sexual, and diploid sexual reproduction, variously under independence of inheritance between loci as well as genetic linkage, fixed fitness and varying fitness conditions.^1^ A summary of some of the results appears in Figure 1.

**Figure 1.**
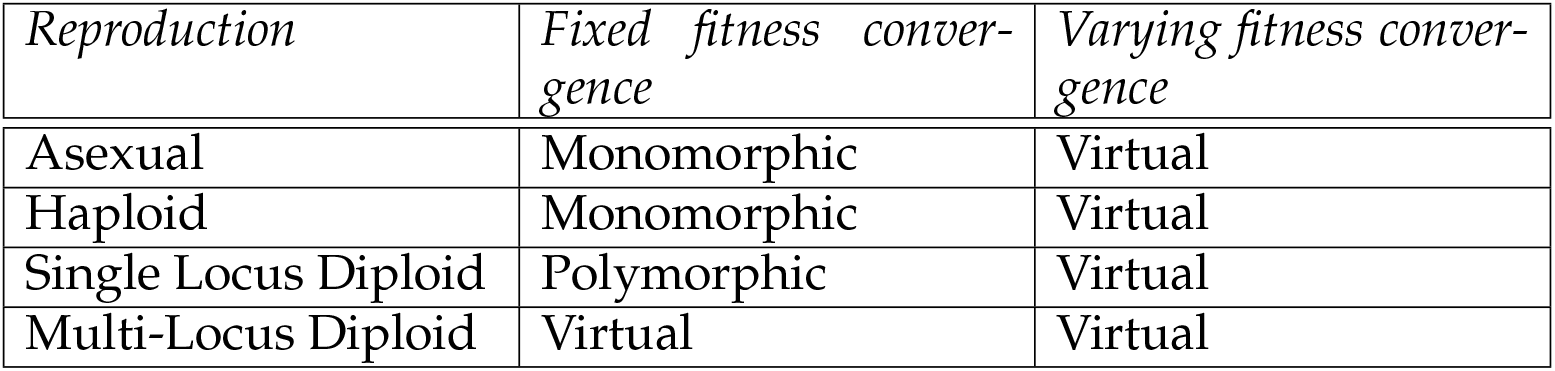
Summary of modes of convergence.

**Figure 2.**
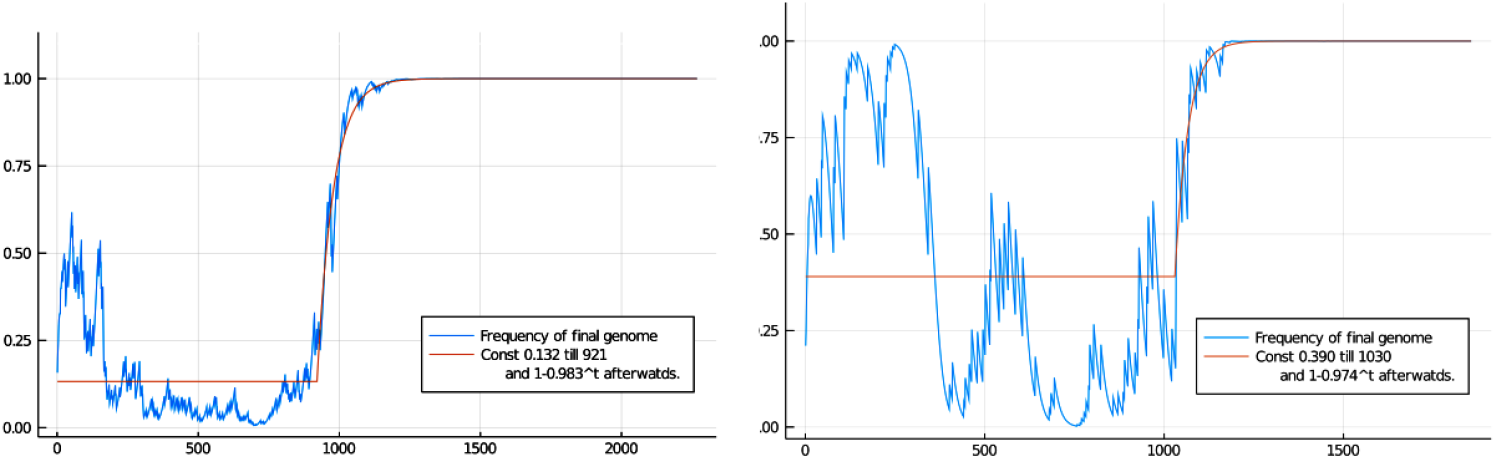
An illustration of the convergence to a monomorphic populations consisting of one genotype under haploid sexual reproduction. In both simulations, the frequency of the final genome in the population over generations is tracked. This frequency appears to drift most of time, then converges at an exponential rate to fixation towards the end, in accordance with Proposition 8.

As can be seen in the summary, all of the reproduction models studied here attain virtual convergence, no matter how wildly environments vary. They attain this by exploiting the regret minimisation aspect of the multplicative weights updating algorithm.

In a sense, it can be said that the reproductive models are opportunistic: when environments are sufficiently well-behaved and they can take advantage of the payoff climbing afforded by the Baum–Eagon inequality, they will do so to converge to Nash equilibrium. If that is not available, then even under the worst possible conditions they will at least attain virtual convergence, minimising regret in hindsight.

### 1.3. Non-arbitrarily Varying Environments

The gap between fixed environments and entirely arbitrarily varying environments is large. The subject of convergence when environments vary in a structural way is explored here only with respect to the asexual replicator model, where we show that convergence to a monomorphic population is guaranteed under ergodically varying environments and under a broader property we introduce that we call one-step-ahead superiority.

There is much scope remaining to be researched in this topic, which we leave to future papers.

## 2. Basic Models and Notation

### 2.1. Simplices

For an integer *m*, Δ^*m*^ denotes the standard finite dimensional simplex over *m* + 1 points. For a finite set Γ, Δ(Γ) denotes the collection of probability mass functions over the elements of Γ. We will denote the subset of Δ(Γ) consisting of distributions with support on one element of *G* alone by Δ_1_(Γ), and the element of Δ_1_(Γ) placing all support on *g* ∈ Γ will be denoted by 1_*g*_.

### 2.2. Potential Games

Let *I* be a finite set of *m* players. Associate with each player *i* a finite set of actions *A_i_*. Denote *A* = *A*_1_ × … × *A_m_*, and the cross product of all action sets except from *i* by *A*_−*i*_. A game is defined by a payoff function *u* : *A* → ℝ^*m*^. The projection of the payoff function to the payoff of player *i* is denoted *u_i_*(*a_i_, a*_−*i*_). Payoff functions extend in the obvious multi-linear manner to payoff functions of mixed strategies.

An identical interests game is a game satisfying the property that *u_i_*(*a*) = *u_j_*(*a*) for each *a* ∈ *A* and each *i, j* ∈ *I*. A potential game is game with a potential function Φ : *A* → ℝ satisfying for all *a*_−*i*_ ∈ A_−*I*_ and all 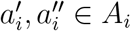

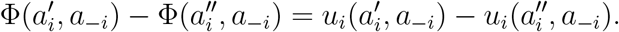

An ordinal potential game is game with a potential function Φ : *A* → ℝ satisfying for all *a*_−*i*_ ∈ *A*_−*i*_ and all 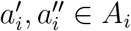

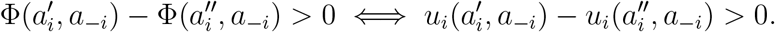

Every identical interests game is a potential game and every potential game is an ordinal potential game.

### 2.3. The Discrete Replicator Equation

Much of the background material for the population genetics models here is from [Bürger, 2000] and [Edwards, 2000].

Time is discrete and denoted by positive integers *t*. Let *f* ^*t*^ : Δ^*m*^ → ℝ^*m*^ be given for each time *t*, with 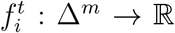 for each 1 ≤ *i* ≤ *m* being the standard coordinate projection of *f ^t^*.

The mean value function associated with *f ^t^*, denoted 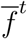, maps *θ* ∈ Δ^*m*^ to ℝ by

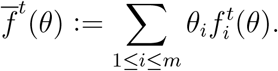

The discrete replicator equation is then the recursive mapping from Δ^*m*^ to Δ^*m*^ defined by

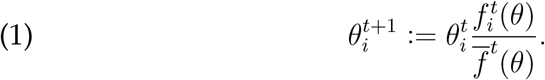

### 2.4. Alleles and Genotypes

The model assumptions which will be maintained throughout are that populations are infinite (i.e., only proportions of genotypes and alleles in the population are of interest, not absolute numbers), that generations are discrete and non-overlapping, that selection occurs but not mutation or migration, and that stochastic genetic drift over time does not occur.

We assume that each gentoype is composed of *m* genetic loci. Each locus *i* is associated with a set of alleles 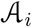 composed of *k* + 1 alleles. A *genotype* is then formally a string *g* = *a*_1_*a*_2_ … *a_m_*, such that 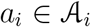 for each 1 ≤ *i* ≤ *m*. Denote the collection of all possible genotypes by Γ.

At each time *t* there is an adult population Π^*t*^ composed of individuals, each of which bears a genotype *g* ∈ *G*. The sub-population of individuals bearing genotype *g* at time *t* is denoted 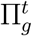.

The adults in the population at time *t* in reproduce (asexually, haploid sexually, or diploid sexually, depending on the particular model being studied). After the adults in population Π^*t*^ reproduce, an offspring population Ω^*t*^ comes into existence. The sub-population of individuals bearing genotype *g* at time *t* is denoted 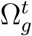

Denote by 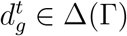 the weight or relative proportion of genotype *g* at time *t* in the offspring population, i.e. the proportion of the set 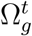 in Ω^*t*^. At the beginning of period *t* + 1, the adult population Π^*t*^ dies, and as the individuals in Ω^*t*^ attain maturity they form the adult population Π*t*+1.

A selection fitness value 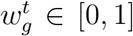 is associated with each genotype *g* at each time *t*. This is interpreted as the probability that an offspring individual bearing genotype *g* in population Ω^*t*^ will survive and attain reproductive maturity as an adult in population Π^*t*+1^.

### 2.5. Asexual (Clonal) Reproduction Model

In this model, at each time *t* each individual in 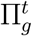 produces *ζ* offspring (where *ζ* is a positive integer), each of whom bears the same genotype *g* as its parent. The offspring thus produced in population Ω^*t*^ then mature into the adults in population Π^*t*+1^, subject to selection as determined by 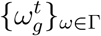

The relevant state space of the dynamic is the simplex Δ(Γ). The *mean fitness* at time *t* is

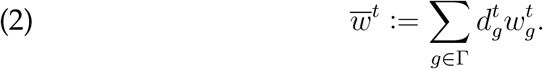

In models in which 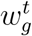 is constant over time we may suppress the denotation and simply write *w_g_*, and hence 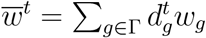

The dynamics of asexual reproduction are governed by the asexual replicator equation as the equation of motion,

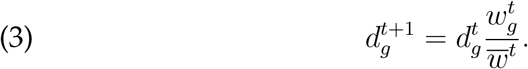

This follows the schema of Equation (1), with the fitness 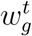 in the role of *f_i_* and 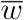 corresponding to 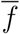.

It will sometimes be convenient to express Equation (3) generically as

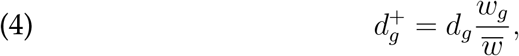

suppressing reference to *t* when its value is clear from context.

### 2.6. Haploid Sexual Reproduction Model

This model will be central to much of the paper, hence we present its assumptions here in some detail. We suppose a monoecious sexually reproducing haploid population, with panmictic mating occuring in pairs. Initially it will be supposed that there is no linkage between loci, i.e., each offspring at each locus bears the allele of one of the parents at the corresponding locus with equal probability. This assumption will be relaxed subsequently.

We will sometimes denote allele *j* at locus *i* by *a_ij_*, where 1 ≤ *i* ≤ *m* and 1 ≤ *j* ≤ *k* + 1, when convenient without confusion by context. When it is important to distinguish the *j*-th allele in locus *i* from the *j*-th allele at locus *i*′, we will explicitly write 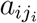. Let 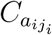 denote the collection of all possible genotypes that contain 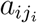 at the slot for locus *i*. Write 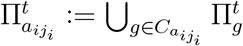 and 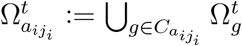.

Denote by 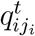 the *allelic frequency* of allele 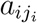 at locus *i* at time *t* in population Ω^*t*^, i.e., 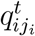 is the proportion of 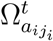 in Ω^*t*^. Call 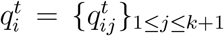 the *allelic frequency distribution* of locus *i* at *t*; this is an element of a *k*-simplex, which we will denote 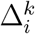. The relevant phase space for studying the evolutionary dynamic is then a polytope composed of an *m*-cross product of simplices:

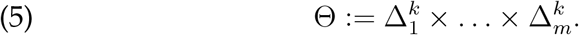

The topology for studying convergence is the product topology of the simplices regarded as manifolds.

Going from Δ(Γ) to Θ is always possible, since we defined 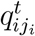 as the proportion of 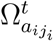 in Ω^*t*^ for each 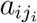. Denote the mapping thus defined by *ρ* :Δ(Γ) → Θ.

For 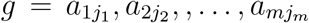 and an allelic frequency distribution *q^t^* ∈ Θ denote

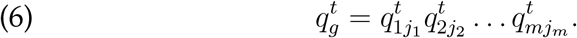

If 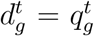 for all *g*, a population is said to be in linkage equilibrium. When linkage equilibrium obtains, the inverse mapping *ρ*^−1^ : Θ → Δ(Γ) is well-defined by applying Equation (6). When we make use of this inverse mapping, given *x* ∈ Θ we will write [*ρ*^−1^(*x*)]_*g*_ to stand for the *g*-th component of *ρ*^−1^(*x*) ∈ Δ(Γ).

The *marginal fitness* of allele *a_ij_* at time *t* is defined as

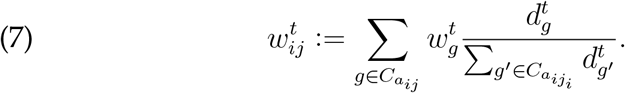

From the collection 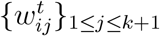 we furthermore can calculate the *mean payoff* for locus *i*, which is 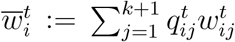 But this yields nothing new, because 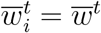 of Equation (2) for all loci *i*.

The dynamic in this model is the *haploid sexual replicator* which can be shown to be

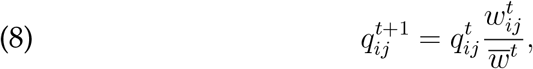

and applies at every allele *j* of every locus *i*. This clearly follows the schema of Equation (1) with 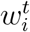 here in the role of *f ^t^* in Equation (1) and 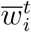 as 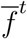. As before, it will sometimes be convenient to express Equation (8) generically as

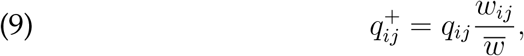

suppressing reference to *t* when its value is clear from context.

The haploid sexual replicator dynamic maps points in Θ to points in Θ, and hence also maps points in 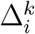 to points in 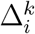 under the projection from Θ to 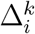.

#### 2.6.1 Haploid Reproduction as an Identical Interests Game

In the fixed fitness case, the collection of fitness values {*w_g_*}_*g*∈Γ_ can be regarded as defining an identical interests game between the loci. H ere we essentially rewrite some of the previous sections in notation that is more familiar from the game theory literature, and unite the analysis of haploid dynamics with that of the dynamics of player strategies in repeated identical interests games when the players implement the haploid sexual replicator equation, Equation (8), in updating their strategies.

From the fixed fitness haploid model with polytope Θ of alleles at the various loci, define an identical interest game *W* _Θ_ as follows. Each locus *i* becomes a player *i*. The set of alleles of locus *i* becomes the set of pure actions *A_i_* of player *i*. For each profile of pure actions (*a*_1_, …, *a_m_*) ∈ *A*_1_ ×…×*A_m_*, the payoff *w_i_*(*a*_1_, …, *a_m_*) = *w*(*a*_1_, …, *a_m_*) to each player *i* is identical and defined to be

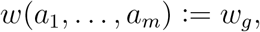

where *g* = *a*_1_ … *a_m_* is the genotype defined by 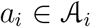 for each *i* and *w_g_* is fitness payoff to genotyper *g*.

Here 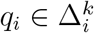, which previously denoted the distribution of alleles in locus *i*, is interpreted as a mixed strategy. The mean fitness 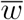 is interpreted as 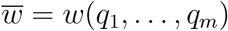, the expected payoff (to each player in the game) when each player/locus *i* plays mixed strategy *q_i_*. The expected payoff/ mean fitness 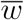 plays the role of the potential function in the identical interests game *W*_Θ_.

Every potential game (and hence every identical interests game) admits at least one pure strategy Nash equilibrium, namely the pure strategy profile yielding the highest potential payoff. The set of all pure Nash equilibria is the set of local maxima of the potential. Denote this set of pure Nash equilibria of *W*_Θ_ by 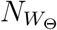.

Each 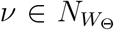 is by definition a profile of alleles (*a*_1_, …, *a_m_*), one from each locus. Hence it is naturally associated with a particular genotype that we will denote *g_*v*_* ∈ Γ.

Note that if the set of mixed strategy profiles is restricted to a subset Θ′ ⊂ Θ, a different identical interests game *W*_Θ′_ is induced. The set of pure Nash equilibria of *W*_Θ′_ may differ from the set of pure Nash equilibria of *W*_Θ_.

We may write 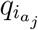 as a synonym for *q_ij_* when *q_i_* is the mixed strategy of *i*. We can write *w*(*p*; *q*_−*i*_) for the expected payoff when locus *i* plays mixed strategy *p* while all the other loci play mixed strategy *q*_−*i*_. In a special case this notation becomes *w*(*a_j_* ; *q*_−*i*_), standing for the expected payoff when pure action/allele 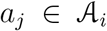 is chosen at locus *i* while all the other loci play mixed strategy *q*_−*i*_; this is none other than the game interpretation of the marginal fitness of allele 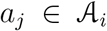, which was above written as *w_ij_*. Then for each *i*,

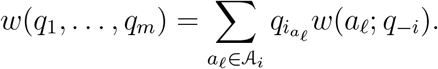

### 2.7. Diploid Sexual Reproduction Model

#### 2.7.1. One Locus

In the single locus diploid model, with a set of alleles 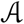, one needs to keep track of pairs of alleles, 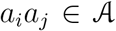, which constitute the genotypes. We suppose no position effects and hence do not distinguish between *a_i_a_j_* and *a_j_a_i_*. Random mating is also assumed, hence Hardy–Weinberg ratios hold during the mating phase (with selection then moving the adult population away from the Hardy–Weinberg ratios).

Label the frequency of allele *a_i_* at time *t* by 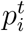 and the frequency of genotype *a_i_a_j_* by 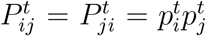. Denote the fitness of genotype *a_i_a_j_* by 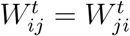, and the population mean fitness by

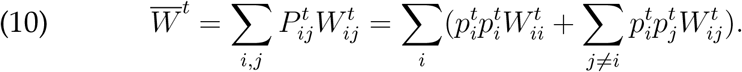

Define the marginal fitness of allele *a_i_* as

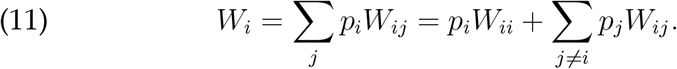

With all the preliminaries in place, the dynamic is once again defined by a straight-forward replicator as the equation of motion

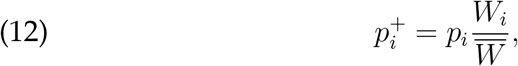

for each allele.

#### 2.7.2. Multiple Loci

The multi-locus diploid model is complicated to describe; we omit most details and present only the minimal notation needed for our purposes here.

As before we suppose that there are *m* loci with *k* + 1 alleles per locus. The state space is 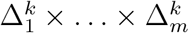 and trajectories are elements 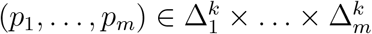.

Within each locus *i*, as in the single locus model, the alleles are between themselves playing at each time period a symmetric potential game with a fitness 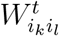 assigned to each pairing 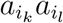, where 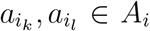. However, 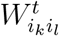 is now a function not only of 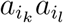 but of the entire profile *p*_−*i*_ of the allelic distributions of the other loci.

The standard analysis in the literature tracks the distribution of gametes (where each gamete is one possible haploid half of a diploid genotype). Each gamete *g* can be assigned a marginal fitness *W_g_* as a function of the fitnesses of the pairings at each locus and the allelic frequency, and from this the mean fitness 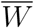 of the population is calculated. Denoting the frequency of gamete *g* by *r_g_*, one can derive a recursion formula that is reminiscent of but not identical to the replicator equation

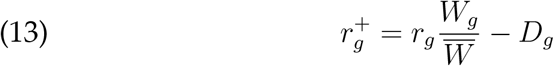

where *D_g_* is the linkage disequilibrium for *g*. The existence of the disequilibrium term *D_g_* means that the diploid multi-locus dynamic is not a replicator dynamic, making the analysis of this dynamic different from all the other models studied in this paper.

## 3. Multiplicative Weights, Potential Games, and Virtual Convergence

### 3.1. Regret Minimisation

The objective of many of the on-line learning algorithms developed in the literature in recent years is the attainment of regret minimisation. Let 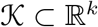 be non-empty, bounded, compact, and convex. At each iteration time *t* algorithm 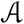 selects an element 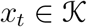, while a concave function 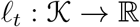 is revealed.

The goal of the algorithm is to minimise the average regret over any *n* rounds, defined as

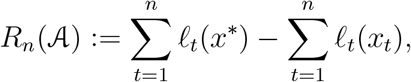

where 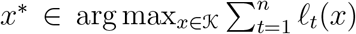. In other words, the objective is to have minimal regret relative to having selected the best possible 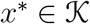 from the start and playing *x*\* in a fixed manner at every time period.

An algorithm implements asymptotic regret minimisation if its regret is sub-linear, i.e., 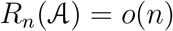 as *n* → ∞. When this holds

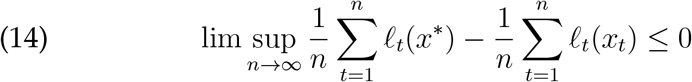

where *x*\* is the element of 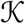 with the optimal average payoff.^2^ In other words, the average regret converges to zero in the limit and the payoff of the algorithm approaches that of having selected the optimal in hindsight *x*\* from the start and monotonically selecting only that point at every iteration.

### 3.2. Multiplicative Weights Update Algorithm

The multiplicative weights update algorithm comes in two flavours: a polynomial and exponential version. In the polynomial version, *d* ∈ Δ^*k*^ is mapped to *d*^+^ ∈ Δ^*k*^, conditional on receipt of a given tuple of real numbers (*ℓ*_1_, …, *ℓ_k_*), by

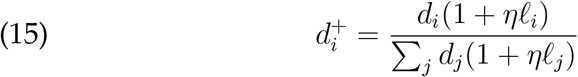

for some *η* > 0. Dividing the numerator and denominator of Equation (15) by *η* changes nothing, hence Equation (15) can be equivalently expressed as

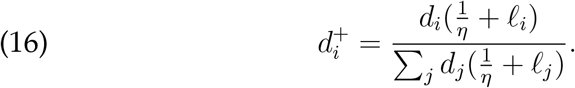

In the special case in which *η* → ∞, sometimes called the parameter-free version of the algorithm (cf. [Meir and Parkes, 2015]), Equation (16) becomes

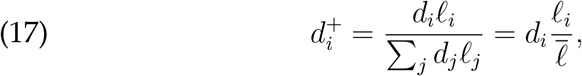

which is exactly the replicator equation.

In its exponential version the multiplicative weights update algorithm, also known as the Hedge algorithm ([Freund and Schapire, 1997])), maps *d* ∈ Δ^*k*^ to *d*^+^ ∈ Δ^*k*^, conditional on receipt of a given tuple of real numbers (*ℓ*_1_, …, *ℓ_k_*), by

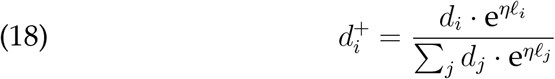

for some *η* > 0.

The replicator equation can also be shown to be a special case of the exponential algorithm ([Edhan et al., 2017]) as expressed in Equation (1). The key is to register not the fitness payoffs a t each time period but the logarithms of the fitnesses: given a fitness tuple *f* = (*f*_1_, …, *f_m_*), form the tuple (*ℓ*_1_, …, *ℓ_m_*) by setting 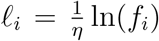. Then apply Equation (18):

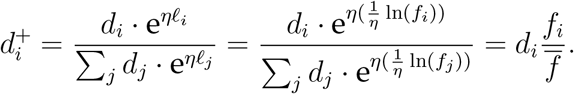

It is well known in the literature that the multiplicative weights update algorithm attains regret minimisation. In the genetic context studied here, this translates into attaining asymptotic average growth rates equal to that of having selected the optimal-in-hindsight genotype *g*\* from the start and hypothetically running history again with a population consisting solely of *g*\* at every time period.

Furthermore, since the haploid sexual reproductive strategy can be interpreted as an implementation of the replicator independently in each locus, the interpretation of the replicator as an instantiation of the multiplicative weights updating algorithm is applicable in several of the models in this paper, beyond the asexual model.

Several papers studying the applicability of multiplicative weights updating algorithms to evolutionary models have been published in recent years. A brief list of such papers includes [Livnat et al., 2008, Chastain et al., 2013, Chastain et al., 2014, Meir and Parkes, 2015].

### 3.3. Multiplicative Weights and Baum–Eagon

It is instructive to compare the multiplicative weights updating algorithm, especially in its parameter-free version:

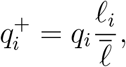

and Baum–Eagon updating (as in Equation (24))

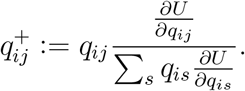

From the perspective of each Δ^*i*^, the Baum–Eagon updating is a special case of the multiplicative weights updating algorithm in which the payoff *ℓ_ij_* is given as the partial derivative of a potential function *U* with respect to *q_ij_*. This perspective will have a significant role here, as many of the dynamics that will be studied benefit both from the monotonic potential increase afforded by the Baum–Eagon Theorem and the regret-minimisation given by the multiplicative weights algorithm aspect.

### 3.4. Convergence in Potential Games

The content of the following theorem is technically equivalent to a theorem in [Palaiopanos et al., 2017] (see also [Panageas et al., 2019]), which is expressed and proved there in the context of congestion games. We present it here with a full proof for two reasons: a) an independent proof for potential games is of value; b) the proof here can readily be understood in the context of reproductive strategies, such as haploid sexual reproduction, given the interpretation of such strategies as implementing the multiplicative weights updating algorithm, as described in Section 3.2, in the context of a potential game between loci, with alleles in the role of pure actions, as described in Section 2.6.1.

#### Theorem 1.

*Suppose that each of a finite set of players playing a potential game implements the polynomial multiplicative weights update algorithm at discrete time periods to update his mixed strategy, starting from a mixed strategy of full support*.

*Then the strategy profile of the players will converge to a fixed point that is a Nash equilibrium*.

Theorem 1 is a stronger result than may appear at first glance, because there is no explicit coordinating element between the players that is assumed. To see why this may be surprising, consider the following extremely simple 2 × 2 game, which is an identical interests game (and hence a potential game):

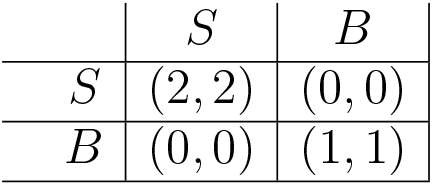

One may interpret this as a coordination game between a couple, who wish to meet. If they are both at the symphony hall (action profile (*S, S*)) they each receive a payoff of 2; if they are both at the beach (action profile (*B, B*)) they each receive a payoff of 1 ; otherwise they fail to meet and receive zero payoff. Suppose that both players simultaneously implement a simple-minded best reply strategy, beginning at action profile (*S, B*). Then in the next time period, the action profile will be (*B, S*), followed by (*S, B*) etc. Lacking a coordinating element, no convergence to a fixed point is attained.

In contrast, Theorem 1 does guarantee convergence under the multiplicative weights update algorithm, even though there is no coordination between the players and each player updates his or her mixed strategy from one time period to the next entirely independently of the other players. It is as if coordination is attained ‘for free’. This result is attained by virtue of the Baum–Eagon Theorem, which underlies the proof of the theorem and guarantees that, despite the lack of coordination, a monotonic climb up the potential of the game ensues at each time period.

### 3.5. Virtual Convergence

Let 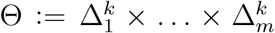 be a polytope, with *T* : Θ → Θ a transformation.

We will say that the dynamic defined by *T converges polymorphically* if for each initial point *x* ∈ Θ there is a point *y* ∈ Θ such that lim_*n*→∞_ *T^n^*(*x*) = *y*. In the special case that for each *x* the limit *y* = lim_*n*→∞_ *T^n^*(*x*) = (*q*_1_, …, *q_m_*) satisfies the condition that *q _i_* is a point distribution in 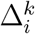 for each 1 *i m*, the dynamic *converges monomorphically*.

Suppose now that a linear fitness function *ℓ_t_* : Θ → ℝ is revealed for each time *t*. For an initial point *x* ∈ Θ, denote *x_n_* := *T^n^*(*x*), with *T* ^0^(*x*) = *x*. We will say that the dynamic defined by *T virtually converges polymorphically* if for any sequence *ℓ*_1_*, ℓ*_2_, … of payoffs and any initial point *x* ∈ Θ, there is a point *y*\* ∈ Θ such that

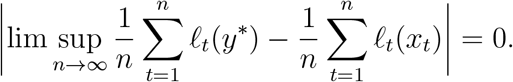

In the special case that virtual convergence is to a *y*\* that is a point distribution in 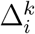 for each 1 ≤ *i* ≤ *m*, we say that *virtual monomorphic convergence* obtains.

## 4. Asexual (Clonal) Reproduction

The dynamics of frequency independent asexual reproduction without mutation is perhaps the simplest of evolutionary dynamics – essentially ‘bacteria in a petri dish’. Despite the apparent simplicity, there is much to be said here that will also have implications for the analysis presented in later sections.

### 4.1. Fixed Fitness

We suppose here a fixed fitness value *w_g_* for each genotype at each time period, generically with a genotype *g*\* ∈ Γ whose fitness *w_g*_* is maximal amongst the genotypes. There are several ways to analyse this; in the spirit of this paper, we may regard this dynamic as a single-player potential game. In this interpretation, there is one player whose mixed strategy at time *t* is a probability measure *d^t^* ∈ Δ(Γ) over the genotypes in Γ. The expected payoff is 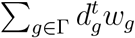. Theorem 1 then implies convergence to a fixed point in Δ(Γ).

Alternatively, we may directly apply the Baum–Eagon Theorem. The dynamics are governed by the asexual replicator equation,

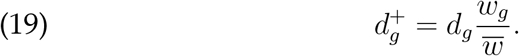

Since 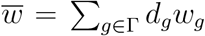, it follows that 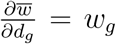, hence Equation (19) is an application of the Baum–Eagon transformation as expressed in Equation (24).

Denote by *T*_0_ : Δ(Γ) → Δ(Γ) the transformation that defines *d*^+^ = *T*_0_(*d*) by mapping *d_g_* to 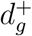 for each *g* according to Equation (19). Since the population mean fitness is increasing montonically, 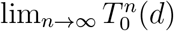 for any starting distribution *d* ∈ Δ(Γ) converges to a point in Δ_1_(Γ), i.e., a fixed point that is a point distribution, since the only fixed points of Equation (19) are point distributions. All the weight is asymptotically on 1_*g*\*_, where *g*\* is the genotype of maximal fitness.

This implies that the interior of the simplex Δ(Γ) forms a global exponentially stable basin of attraction. If, in contrast, the initial point lies within a strict subface *F* ⊂ Δ(Γ), then the convergence will again be to a monomorphic population whose genotype is the genotype of maximal fitness within *F*. This will clearly be sub-optimal if *g*\* ∉ *F*.

### 4.2. Temporally Varying Fitness

The fixed fitness setting of asexual reproduction is the simplest evolutionary model, yielding perhaps the strongest result that can be expected, of monotonic and rapid fitness increase towards convergent fixation to the globally optimal genotype. This satisfactory result, however, may not necessarily obtain if fitnesses are no longer fixed in time.

In a temporally varying fitness model, we suppose that there is a collection of possible environments Ω, such that each *ω* ∈ Ω determines a fitness landscape such that each genotype *g* is assigned a fitness value *w_g_*(*ω*) under *ω*. At each time *t* one environment *ω* from Ω is selected, with the payoff to the genotypes registered in accordance to the fitness landscape of that environment.

A simple hill climbing dynamic cannot be applicable here because there is a different ‘hill’ (i.e., fitness gradient derived from the environment) at each time period; the trajectory under the transformation *T*_0_ will no longer be monotonically increasing in mean fitness. Despite this, the replicator algorithm does an excellent job at learning, even under conditions of temporally varying fitness. This can be seen in several ways.

Consider first a discrete i.i.d. model in which there is a probability measure *μ* over Ω determining the selection of the environment at each time period, repeated indefinitely. This determines for each genotype *g* an expected fitness payoff under *μ*. An optimal population will (generically) be composed of the genotype with maximal expected fitness payoff, and the replicator reliably identifies this genotype. More generally:

#### Proposition 1.

*Let 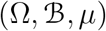 be a probability space over a collection Ω of environments. For each genotype g ∈ Γ, define a random variable w_g_(ω) ∈ [0, 1], interpreted as the fitness of g under environment ω ∈ Ω, from which the expected fitness is given as E (w_g_ | μ) = ∫_Ω_ w_g_(ω) dμ(ω)*.

*Let S : Ω → Ω be a stationary and ergodic transformation defining a stochastic process for each g by 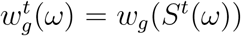). Then under the asexual replicator dynamic, with probability one the population asymptotically converges to a monomorphic population consisting of the genotype with maximal expected fitness*.

Proposition 1 indicates that when there is sufficient structure to the stochastic process of the varying environments, at least as expressed in stationary ergodicity (which include i.i.d. as a special case), the replicator dynamic will be able to extract the information inherent in the process to identify the optimal genotype and converge to that genotype, from any initial population state (that at least minimally includes the optimal genotype).

From here one can ask what happens when the stochastic process of varying environments can be any process at all. It is not difficult to conjure examples of temporally varying environments that do not admit convergence to a single genotype. For example, let

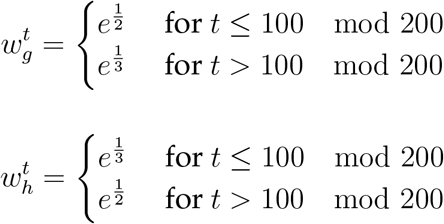

Then clearly both 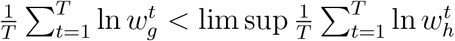 and 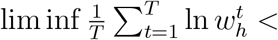 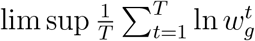. When one genotype is strong the other is weak, each temporarily overtaking the other only to fall back later.

Nevertheless, it is possible to extend Proposition 1 to much more general environments using the notion of *one-step-ahead expected log-fitness*. The one-step ahead expected log-fitness is the expected log-fitness of a generation conditional on the past generations.

#### Definition 2.

*Let 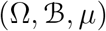 be a probability space over a collection Ω of environments and let (ψ_t_)_t≥1_ be a stochastic process of environments relative to 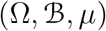. For each genotype g Γ, define a process by 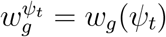, interpreted as the fitness of g under environmental processψ _t_. Let 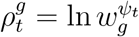 denote the log-fitness; assume that 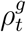 is always bounded*.

*We will call 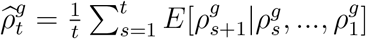 the average one-step-ahead expected log-fitness of g at t*.

#### Definition 3.

*A gentoype g is asymptotically one-step-ahead superior on average if 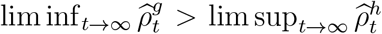 with probability one for all genotypes h ∈ Γ with h ≠ g*.

#### Theorem 2.

*If a genotype g ∈ Γ is asymptotically one-step-ahead superior on average then, under the asexual replicator dynamic, with probability one the population asymptotically converges to a monomorphic population consisting of the genotype g*.

It is worthwhile noting here that in the case of an ergodic environment, 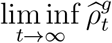 and 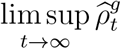 are one and the same and equal to the constant *E*[ln *w_g_*] almost surely. Thus the sufficient condition established in Theorem 2, namely asymptotic one-step-ahead superiority, is reduced to *E*[ln *w_g_*] > *E*[ln *w_h_*].

The statement of Theorem 2 supposes that genotype *g* is asymptotically one-step-ahead superior on average with probability one with respect to all environments. Suppose instead that a genotype *g* is is asymptotically one-step-ahead superior on average only with respect to a subset 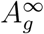 of the collection of environments. Then we obtain the following corollary.

#### Corollary 4.

*If 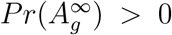, where 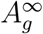 is the set of environments in which genotype g is is asymptotically one-step-ahead superior on average, then under the asexual replicator dynamic, with probability one in 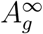 the population asymptotically converges to a monomorphic population consisting of the genotype g*.

Algorithms such as the multiplicative weights and mirror ascent algorithms have been developed in the computer science literature in recent years for the sake of optimisation under conditions of no statistical structure. The replicator dynamic, it turns out, exploits the results afforded by these algorithms.

#### Theorem 3.

*Under the replicator dynamic, for any arbitrary temporally varying fitness there is an optimal-in-hindsight genotype g* such that for any initial point in the interior of the simplex, asexual reproduction virtually converges monmorphically to g\**.

In summary, we interpret the results of this section from a learning perspective: the objective is to learn which genotype is best fit for the environment process, via the algorithmic tool of the replicator.

When the environment is fixed, the replicator homes in on the objectively fittest genotype. When the environment process is sufficiently structured, as in a stationary ergodic process, the replicator makes use of time averaging to identify a winning genotype. Failing that, in the worst case in which there is insufficient structure for predictive learning, the replicator still manages to extract information, by application of regret minimisation via the multiplicative weights updating algorithm; virtual convergence occurs in the sense that one can imagine a population which from the start consisted of only the optimal-in-hindsight genotype and attaining the same asymptotic average growth rate as actually attained.

## 5. Haploid Sexual Reproduction

### 5.1. Fixed Fitness

In this section the population will be presumed to reproduce via haploid sexual reproduction under a fitness landscape {*w_g_*}_*g*∈Γ_ that is fixed throughout time.

#### 5.1.1. Under Linkage Equilibrium

Under linkage equilibrium, in population 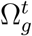 the equation 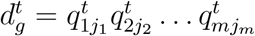 holds for each genotype 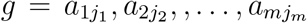 As 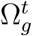 matures into 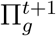, selection applies such that linkage equilibrium does *not* hold for 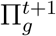; however, by assumption random mating between the reproducing adults in 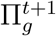 immediately restores linkage equilibrium in the next offspring generation 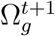.

One advantage of working with an assumption of linkage equilibrium is that we may identify in a bijective manner a point in the allelic frequency space Θ and a corresponding point in the genotypic frequency space Δ(Γ). We shall freely do so in this section as follows.

Recalling the haploid sexual replicator,

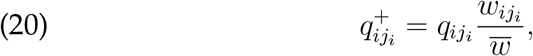

define *τ* : Θ → Θ to be the transformation given by the mapping of *q_ij_* to 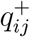 for each locus *i* and allele *j* in *i*. Exploiting the linkage equilibrium assumption, define a transformation *T*_1_ : Δ(Γ) → Δ(Γ) by

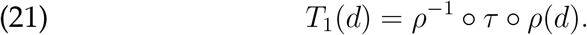

Abusing terminology, we will call both *τ* and *T*_1_ haploid sexual replicator transformations. This enables us to analyse the dynamics equally well under either *T*_1_ or *τ* ; both define discrete dynamical systems determining trajectory paths in Δ(Γ) or in Θ, respectively.

As in the asexual case, the Baum–Eagon inequality applies (see a similar argument in [Novak and Barton, 2017]). The domain is the polytope Θ as defined in Equation (5).

##### Lemma 5.

*The haploid sexual replicator transformation (under linkage equilibrium and without genetic linkage between loci) satisfies the Baum–Eagon inequality, with mean fitness 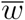 as a Lyapunov function*.

It follows that the population will asymptotically converge to a fixed point of the dynamics defined by the transformation *T*_1_ along paths of monotonically increasing mean fitness.

From here it will be convenient to continue the analysis from the equivalent perspective of the identical interests game interpretation.

##### Theorem 4.

*Under haploid recombinative sexual reproduction (under linkage equilibrium and without genetic linkage between loci), trajectories almost always increase mean fitness monotonically*.

*Beginning from almost any interior point of Δ(Γ) the haploid sexual replicator dynamic converges asymptotically to a monomorphic population in which each individual bears a genotype g_*v*_ from the set 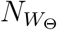 of pure Nash equilibria of the associated potential game W_Θ_*.

##### Corollary 6.

*If the initial point of the allelic frequency of the population lies in any face Θ′ of Θ then the dynamic converges asymptotically to a monomorphic population consisting of genotypes from the set of pure Nash equilibria of the associated potential game W′_Θ_*.

There are immediate interesting implications of Theorem 4. One of these is that Δ(Γ) is entirely partitioned into asymptotically stable basins of attraction (deterministically in this model).

##### Theorem 5.

*For each pure Nash equilibrium 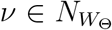, there exists *B_*v*_* ⊂ Δ(Γ) containing 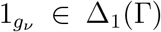 such that starting from any initial point in B_*v*_ the population under the dynamic will converge to a monomorphic population consisting solely of genotype g_*v*_, i.e., 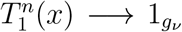 for every *x* ∈ *B*_*v*_. Apart from separatrices between these basins of attraction, which are of negligible measure, the sets in the collection {*B_*v*_*} form a partition of Δ(Γ)*.

Even more than that can be said here. By Theorem 6 of [Baum and Sell, 1968], any transformation of the form defined in Equation (24) increases *U* - homotopically, from which it follows that the haploid sexual replicator transformation *T*_1_ : Δ(Γ) → Δ(Γ) increases 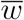-homotopically.

##### Proposition 7.

*Let *S_t_*(*x*) = *tT*_1_(*x*) + (1 − *t*)*x*. For each pure Nash equilibrium 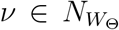, there exists a neighbourhood *H*_*v*_ ⊂ Δ(Γ) of 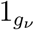 such that *S_t_*(*H_*v*_*) ⊂ *H_*v*_* for 0 < t ≤ 1, and for every 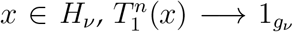. Furthermore, each H_*v*_ has the homotopy type of a disk*.

The significance of the ‘basin of homotopic attraction’ *H_*v*_* of Proposition 7 is that not only does every point *x* ∈ *H_v_* converge to *g_*v*_* under the dynamic, also a small perturbation of around *x* perserves this property. In contrast, around any pure strategy point that is neither a local maximum or a local minimum there are points such that a small perturbation can lead to asymptotic convergence to different fixed points.

Finally:

##### Proposition 8.

*For each pure Nash equilibrium 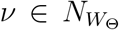, there exists a neighbourhood *E*_*v*_ ⊂ Δ(Γ) 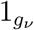 that is an exponentially stable basin of attraction*.

Within the exponentially stable basin of attraction around a Nash equilibrium, the haploid sexual replicator dynamics resembles the asexual replicator dynamics, with exponential convergence to an equilbrium point.

The containment relations are 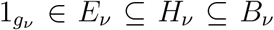. This implies that an observer following a trajectory starting in *B_*v*_* far from *g_*v*_* will likely initially see a slow and moderate increase in mean fitness, with broad polymorphism, for a long time, but once the trajectory enters *E_*v*_* suddenly an extremely fast rise in mean fitness will be registered along with rapid convergence to a monomorphic population.

#### 5.1.2. Under Genetic Linkage Between Loci

Genetic linkage in this section means physical linkage between loci: in the context of haploid reproduction, this means that an offspring zygote might inherit a pair (or more) of alleles from one of the parents as a package, in contrast to an assumption of independent inheritance with probability 0.5 from each parent at each locus.

The term linkage disequilibrium refers to a population genotype distribution that does not equal the cross product of the marginal distribution as reflected in the allelic distribution. We note that in the game theoretic terms that we have been applying throughout to the study of genetic reproduction, linkage equilibrium corresponds to independent strategy selection of each player, while linkage disequilibrium corresponds to dependencies in the selection of strategies.

We initially present an analysis of the haploid sexual replicator under genetic linkage between loci in the case of two loci, for clarity of exposition.

Let *r* ∈ [0, 1] be the recombination rate. Suppose one starts with a point *d* ∈ Δ(Γ) representing the population distribution. Project *d* to Θ via *θ* = *ρ*(*d*). Under the asexual replicator *d* is mapped to *T*_0_(*d*), and under the haploid replicator *θ* is mapped to *τ* (*θ*). Then the replicator equation under recombination rate *r* is

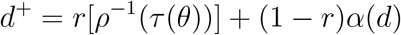

or, using the transformation *T*_1_ defined in Equation (21),

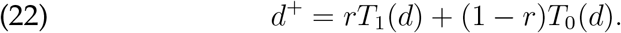

We may denote by *T_r_* the transformation defined by Equation (22) which is consistent with our labelling of *T*_1_ and *T*_0_. When *r* ≠ 1, genetic linkage between the loci occurs.

The recombination rate *r* is intended to describe a situation in which each offspring is produced by sexual recombination with probability *r* and is produced by asexual cloning with probability 1 − *r*. This results in an offspring population, such that within that population, a weight *r* of the offspring is descended from a sexual reproduction event and weight 1 − *r* is descended from an asexual reproduction event.

We may instead consider the following situation, which is mathematically equivalent and more convenient for our purposes: Create two separate copies Π_0_ and Π_1_ of the reproducing population Π, maintaining the genotype frequencies of the original population in each copy, with relative population size proportions 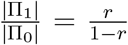. Let Π_0_ reproduce asexually to produce offspring population Ω_0_ and Π_1_ reproduce haploid sexually to produce offspring population Ω_1_, finally combining them into Ω = Ω_0_ ∪ Ω_1_ and regarding the genotypic frequency of Ω.

Slightly more generally, select fraction *r* of the population at random to reproduce by the haploid sexual transformation, with the remaining 1 − *r* of the population reproducing by the asexual transformation. All of these alternatives result in an offspring population with weight *r* descending from a sexual reproduction event and weight 1 − *r* descending from an asexual reproduction event, which is what is relevant.

##### Proposition 9.

*In a population starting at an initial point in linkage equilibrium, under two-locus haploid recombinative sexual reproduction with recombination rate r, trajectories always increase mean fitness monotonically*.

*Beginning from any such interior point of Δ(Γ) the haploid sexual replicator dynamic converges asymptotically to a monomorphic population in which each individual bears a genotype g_*v*_ from the set 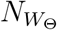 of pure Nash equilibria of the associated potential game W_Θ_*.

In greater generality, suppose that there are *m* loci. Let *λ* be a partition of {1, …, *m*} into *ℓ* ≤ *m* partition elements. An individual will be of *λ*-type if, when reproducing, the genes of that individual undergo physical genetic linkage according to *λ*. In other words, if two *λ*-type individuals *I*_1_ and *I*_2_ mate and produce an offspring *O*, then for each partition element 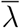 of *λ*, all the alleles in the loci included in 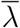 in the genotype of *O* will be identical to either the alleles of 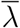 in the genotype of *I*_1_ or the alleles of 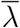 in the genotype of *I*_2_, with equal probability. If the entire population reproduces in this way, denote the resulting transformation from Δ(Γ) to Δ(Γ) by *T_λ_*.

If *λ* is the coarsest partition, consisting of only one partition element, this describes asexual reproduction. For any other partition, *λ*-type reproduction with 1 < *ℓ ≤ m* partition elements reduces to haploid sexual reproduction: simply regard the *ℓ* partition elements as *ℓ* independent loci. If *λ* is the finest partition, in which each locus is its own partition element, this describes haploid sexual reproduction under independence of inheritance at each locus.

Let Λ be the set of Lall partitions of {1, …, *m*}. For each *λ* ∈ Λ let *r*_λ_ ∈ [0, 1], such that Σ_λ∈Λ_ *r*_λ_ = 1. Interpret r_λ_ as the probability that an offspring is produced by physical genetic linkage in accordance with partition *λ*. Mathematically this is equivalent to selecting at random at each generation, for each *λ* ∈ Λ, a fraction *r_λ_* of the population which reproduces by *λ*-type reproduction.

The resulting {*r_λ_*}_*λ*∈Λ_-tuple replicator equation is

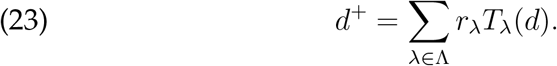

##### Theorem 6.

*In a population starting at an initial point in linkage equilibrium, under m-locus haploid recombinative sexual reproduction with recombination tuple {r_λ_}_λ∈Λ_, trajectories always increase mean fitness monotonically*.

*Beginning from any such interior point of Δ(Γ) the haploid sexual replicator dynamic converges asymptotically to a monomorphic population in which each individual bears a genotype g_*v*_ from the set 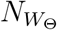 of pure Nash equilibria of the associated potential game W_Θ_*.

The conclusion is that whether or not there is genetic linkage, in haploid sexual reproduction mean fitness increases monotonically and the population always converges to a monomorphic population corresponding to a pure Nash equilibrium of the potential game (this statement also holds true for asexual reproduction, since the equilibrium of maximal mean fitness is itself a pure Nash equilibrium of the potential game). Results similar to those in Theorem 5 and Proposition 7 also attain whether or not there is genetic linkage, with sensitivity to intial conditions as before.

However, although under any genetic linkage structure convergence to some pure Nash equilibrium occurs, the probability of converging to any particular pure Nash equilibrium, starting from the same initial allelic distribution, differs from one linkage structure to another. The same initial point can converge to different equilibria points depending on the linkage structure (as can be seen for example in the extreme case of no recombination, under which convergence will always be to the asexual globally optimal fitness equilibrium.)

#### 5.1.3. External and Internal Environments

Let *ɛ* be a collection of possible environments. Each *e* ∈ *ɛ* determines a fitness landscape in the sense that *e* is identified with an identical interests game *W_e_* played by the loci, relative to a fixed allelic frequency space Θ.

In this section we will make a distinction between what we term the ‘external environment’ and the ‘internal environment’. The external environment is the realisation *e*^1^*, e*^2^, …, *e^t^*, … and the corresponding 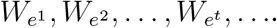 Parallel to this, we may take the perspective of any particular locus *i*. From this perspective, the alleles in locus *i* are amongst themselves implementing an asexual replicator dynamic as follows. Let 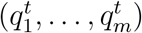 denote the profile of allelic frequencies over time. At time *t*, the identical interests game is 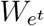, and we may write the time *t* growth rate as the stage *t* game payoff 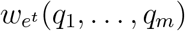 Let 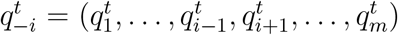 denote the profile of the *m* − 1 loci apart from *i*. Call the sequence 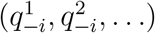 the *internal environment* from the perspective of locus *i*.

Define the *total environment* from the perspective of locus *i* at time *t* to be 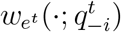, meaning that each choice of *q_i_ ∈ Δ(A_i_)* yields the payoff 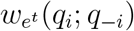. In this way we may reduce the dynamic of each locus to the asexual replicator, with the alleles in locus *i* implementing the asexual dynamic with respect to the total environment from their perspective.

#### 5.1.4. Virtual Convergence

Table 1 presents examples of two fitness matrices with two loci and three alleles per locus. One may interpret these as representing a population that is exposed to two possible environments, one per matrix, where the top is is interpreted as a ‘rainy year’ environment and the bottom one is a ‘drought year’ environment.

**Table 1.** Two examples of fitness matrices for haploid reproduction with two loci and three alleles per locus

| | $a_{21}$ | $a_{22}$ | $a_{23}$ |
| --- | --- | --- | --- |
| $a_{11}$ | $w_{a_{11}, a_{21}} = .40$ | $w_{a_{11}, a_{22}} = .60$ | $w_{a_{11}, a_{23}} = .80$ |
| $a_{12}$ | $w_{a_{12}, a_{21}} = .48$ | $w_{a_{12}, a_{22}} = .55$ | $w_{a_{12}, a_{23}} = .75$ |
| $a_{13}$ | $w_{a_{13}, a_{21}} = .20$ | $w_{a_{13}, a_{22}} = .51$ | $w_{a_{13}, a_{23}} = .70$ |
| $a_{11}$ | $w_{a_{11}, a_{21}} = .40$ | $w_{a_{11}, a_{22}} = .48$ | $w_{a_{11}, a_{23}} = .20$ |
| $a_{12}$ | $w_{a_{12}, a_{21}} = .60$ | $w_{a_{12}, a_{22}} = .55$ | $w_{a_{12}, a_{23}} = .51$ |
| $a_{13}$ | $w_{a_{13}, a_{21}} = .80$ | $w_{a_{13}, a_{22}} = .75$ | $w_{a_{13}, a_{23}} = .70$ |

The top matrix has one pure Nash equilibrium, at (*a*_11_*, a*_23_), and the bottom matrix similarly one at (*a*_13_*, a*_21_). It is possible, however, for an environment realisation to create a trajectory that almost always remains extremely close to points that are not Nash equilibria – even though from the perspective of both matrices there is a repulsion from those points. For example, there can be a trajectory that begins close to (*a*_13_*, a*_21_) and remains there under an alternating realisation, rainy one year and drought the next. This is because although at each time period the single period dynamic pushes away from (*a*_13_*, a*_21_), the directions of push away from that point are nearly opposite, hence the trajectory never wanders far.

Alternatively, it is possible to imagine environment realisations in which very long stretches of one environment bring the population very nearly to convergence to (*a*_11_*, a*_23_), followed by equally long stretches subsequently driving the population very nearly to convergence to (*a*_13_*, a*_21_), repeated periodically in such a way that there is no candidate even for ‘near convergence’. These simple examples indicate that the behaviour under time varying fitness can be complex and highly dependent on initial conditions and random realisations. Nevertheless, we can state the following theorem on virtual convergence. The intuition behind it is that even when the external and internal environments change upredictably, from the perspective of each individual locus the internal alleles are implementing the simple replicator equation, hence each locus experiences virtual convergence.

##### Theorem 7.

*The hapoid sexually reproducing dynamic virtually converges monomorphically under any environment realisation*.

Note that although the result of Theorem 7 guarantees virtual monomorphic convergence under every environment realisation, different realisations can lead to different virtual growth rates. If the external environment process is sufficiently regular, however, then every realisation will lead to the same virtual convergence, even though the corresponding internal environment process may not follow parallel regularity.

##### Proposition 10.

*If the external environment follows a stationary and ergodic stochastic process then every environment realisation leads to the same virtual monomorphic growth rate*.

## 6. Diploid Sexual Reproduction

### 6.1. Single Locus Model

#### 6.1.1. Fixed Fitness

The diploid sexual reproduction in the single locus model is extremely similar to the haploid two locus model, which enables many of the results from haploid sexual reproduction to be carried over almost entirely (and arguably relatively simply, since the parallel is to two loci and not *m* loci) but for one very significant difference: where in the haploid two locus model fitness is represented by a potential matrix with a separate set of alleles for the row player and the column player (corresponding to different alleles in the different loci), in the single locus diploid model the same alleles appear as both row players and column players in the matrix.

The state space is the allelic frequency space Θ = Δ(*A*), with the trajectories in Δ(*A*) recursively following the replicator equation. The fitness landscape^3^ determined by fitness *W_ij_* corresponding to gamete *a_i_a_j_*, denoted here as before by *W*_Θ_ is a symmetric matrix.

Dynamics with respect to symmetric matrices have long been studied in the literature of evolutionary game theory. The parallels are clear: both the diploid single locus and the population dynamic cases can be thought of as a single player game, in which the player selects a mixed strategy (e.g., the ratio of hawks to doves in the population, or the ratio of allele A to allele B), receives an expected payoff, and in the next time period updates the mixed strategy in accordance with a replicator equation. In the evolutionary game theory literature, it is well known that such a dynamic leads to convergence to evolutionarily stable Nash equilibria, which may be either pure or mixed Nash equilibria.

In comparison with the haploid sexual model, the significant new element is, of course, the possibility of convergence to mixed Nash equilibria. One possible evolutionary advantage of maintaining mixed equilibria versus convergence to pure equilibria is that mixed equilibria may be similar to constantly re-balanced portfolios in investment theory; it is well known that re-balanced portfolios (analogous to mixed equilibria) can significantly outperform single stock portfolios (analogous to pure equilibria).

There is one major difference between the dynamics of the diploid sexual reproduction and the population dynamics of a typical evolutionary game theory model. In the fixed fitness/matrix setting, evolutionary game theory models can frequently exhibit cyclic or chaotic trajectories that never converge. In contrast, diploid single locus dynamic trajectories are generically monotonically increase in mean fitness and converge, to either a stable monomorphic population (pure Nash equilibrium) or a stable polymorphic population (mixed Nash equilibrium).

This is because the diploid single locus dynamic follows the Baum–Eagon inequality, montonically increasing in mean fitness. Theorem 1 is not directly applicable here, because of the non-linearity in the payoffs to the players with respect to the payoff matrix. One can instead show directly from the equations of motion that the Baum–Eagon inequality holds. This result is known in the literature (see for example [Edwards, 2000]); we reproduce it here for completeness.

Recall that by Equation (11), *W_i_* = Σ_*i*_ *p_i_W_ij_* and that by Equation (10) 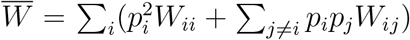. Calculating the partial derivative, 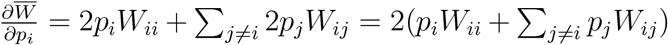 (taking into account the fact that *i* will appear both in *W_ij_* and *W_ji_* for each *j*).

It follows that 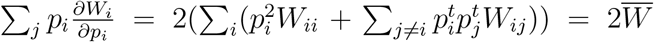.

Hence

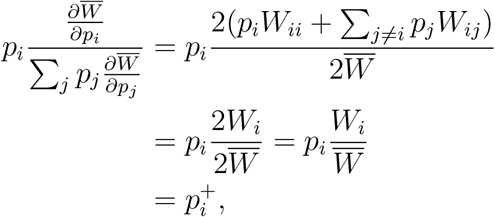

with the last equality following Equation (12). Hence the Baum–Eagon Theorem applies to the dynamic in Δ(*A*) and one concludes that in the single locus diploid sexual reproduction dynamic fitness monotonically increases until a local maximum is attained.

The upshot is that, apart from the possible convergence to stable polymorphism when mixed strategies are the end result of the dynamic, the diploid single locus model parallels the haploid two-locus model in the crucial aspects of monotonic Baum–Eagon mean fitness increase while following a replicator recursion. This enables us to adapt many of the results from the haploid analysis to the diploid model.

##### Theorem 8.

*In the diploid single-locus model, for each ESS equilibrium *v*, there exists B_*v*_ ⊂ Δ(Γ) containing 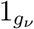 such that starting from any initial point in B_*v*_ the population under the dynamic will converge to a monomorphic population consisting solely of genotype g_*v*_, i.e., 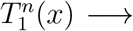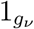 for every *x* ∈ B_*v*_. Apart from separatrices between these basins of attraction, which are of negligible measure, the sets in the collection {B_*v*_} form a partition of Δ(Γ)*.

##### Proposition 11

*Let S_t_*(*x*) = *tT*_1_(*x*)+(1 − *t*)*x. In the diploid single-locus model, for each ESS equilibrium ν, there exists a neighbourhood H_ν_* Δ(Γ) *of* 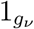 *such that S_t_*(*H_ν_*) ⊂ *H_ν_ for* 0 < *t* ≤ 1, *and for every x* ∈ *H_ν′_* 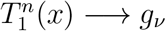. Furthermore, each H_ν_ has the homotopy type of a disk.

#### 6.1.2. Temporally Variable Fitness

The time varying fitness results of the haploid two-locus model similarly carry over to the diploid single-locus model.

##### Theorem 9

*The single-locus diploid sexually reproducing dynamic under varying fitness environments virtually converges polymorphically under any environment realisation*.

### 6.2. Multiple Locus Model

The diploid multi-locus model under linkage equilibrium re-capitulates the diploid single locus model at every locus. Hence the results of the previous section apply without changes.

Many of the tools of the previous sections fail to apply in the diploid multi-locus model under linkage disequilibrium. The main equation of motion, Equation (13), is similar to but not quite a replicator equation. More to the point, the presence of the disequilibrium term causes the dynamic to fail to conform to the Baum–Eagon conditions. Hence the Baum–Eagon theorem, even under fixed fitness, cannot be used to conclude that monotonic fitness increase occurs; indeed, it has long been known that there are examples of fitness landscapes in which diploid populations can exhibit reductions in mean fitness over stretches of time and even periodically cycling trajectories. This leaves no possibility for a general theorem on either monomorphic or polymorphic convergence.

However, virtual convergence (possibly polymorphic), which does not depend on monotonic fitness increase, does obtain here under both fixed and temporally varying fitness. The proof is essentially the same as the proof of haploid virtual convergence (Theorem 7) under temporally varying fitness. As before, we suppose a choice of environment realisation *e*^1^, *e*^2^,… from a set of possible environments, and distinguish between the external environment, represented by such a realisation, and the internal environment perceived by a locus *i*, which is the allelic frequencies of the other loci at any time *t*.

#### Theorem 10

*The multi-locus diploid sexually reproducing dynamic virtually converges polymorphically under any environment realisation*.

## APPENDIX A Baum–Eagon Dynamics

We make use extensive use here of the Baum–Eagon inequality (originally developed for the study of hidden Markov models by use of the Baum–Welch algorithm). The concepts and results in this section are from [Baum and Eagon, 1967] and [Baum and Sell, 1968]. A brief exposition on the Baum–Eagon inequality with applications to population genetics appears in [Edwards, 2000].

## Baum–Eagon Inequality

Let Θ be a polytope given by a cross product of simplices,^4^ i.e., 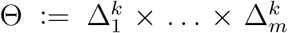 Denote the *j*-th element in the *i*-th simplex by *x_ij_*.

Let *U* (*x_ij_*) be a real-valued polynomial function with non-negative coefficients over the variables {*x_ij_}_i,j_*. Let **x** be a point in the domain Θ. Let *T* (**x**) denote the point of Θ whose *i, j*-th coordinate is given by

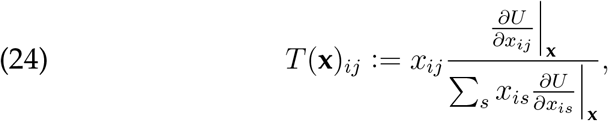

where the denominator is a normalising element.

Then *U* (*T* (**x**)) > *U* (**x**) unless *T* (**x**) = **x**.

It is possible to give the Baum–Eagon inequality an interesting gradient interpretation. Fix *i*, i.e., concentrate on the *i*-th simplex, with each element of 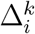 denoted as a tuple **x***i* := (*x_i_*_1_*, x_i_*_2_,…, *x_ik_*). From the perspective of 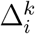 may be considered to be a ‘potential’ function, involving (*x_i_*_1_*, x_i_*_2_,…,*x_ik_*) and other parameters.

Consider a Euclidean gradient vector derived from the potential in this perspective, that is, 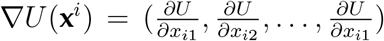. Then the transformation of Equation (24) can be considered as mapping **x**^*i*^ to Δ*U* (**x**^*i*^) · **x**^*i*^ for each *i* separately, followed by projection to the simplex by way of the normalisation. In a sense, the Baum–Eagon dynamic is an application of a form of ‘gradient hill climbing’, locally within each simplex of the polytope Θ, that taken together ensures a global climb.

## APPENDIX B PROOFS

### Proof of Proposition 1.

This is a straight-forward application of Birkhoff’s ergodic theorem. Again we register the log fitness. By the ergodic theorem, for each *g*, 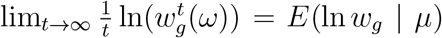 with probability one. Hence the genotype *g*\* with the greatest expected log fitness (which is also the one with the greatest expected fitness) dominates, as it grows at the fastest average rate.

### Proof of Theorem 2.

Consider the field 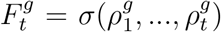 and the random variable

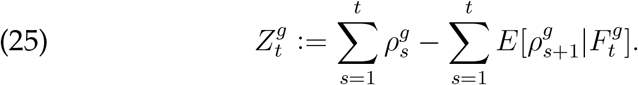

This is a martingale since

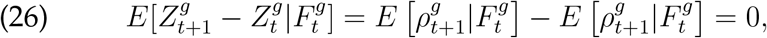

and it is bounded by assumption. Thus, by the Azuma-Hoeffding inequality:

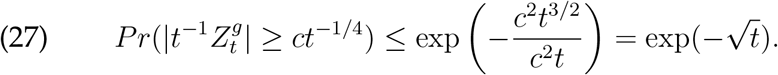

Let *E_t_* be the event that 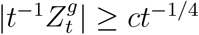. Then we have shown that

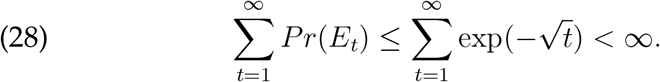

Using the Borel-Cantelli Lemma we deduce

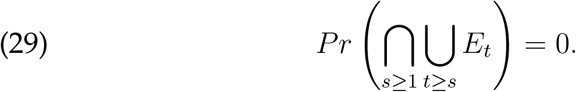

Notice that

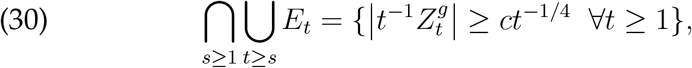

implying that almost surely 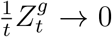. By the definition of 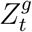 we deduce that almost surely

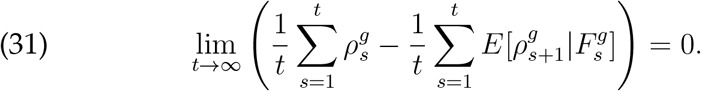

Thus, by adding 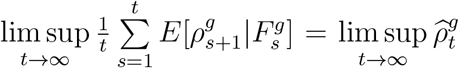 to both sides of the equals sign in Equation (31), we obtain that almost surely

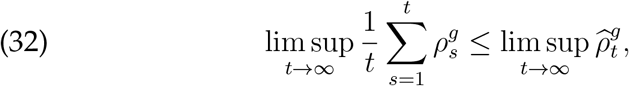

and by an entirely similar argument

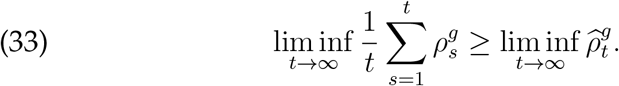

Next, recall that by assumption *g* is asymptotically one-step-ahead superior on average, meaning that by definition lim inf 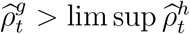 for all genotypes *h* ≠ *g*. Combining this with the inequalities in (32) and (33), which holds for every genotype, one obtains that for every genotype *h*

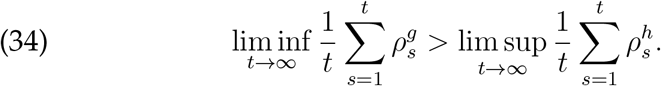

This is sufficient to deduce the statement of the theorem.

### Proof of Corollary 4.

As 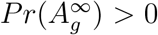 by assumption, we can con-sider the process 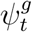 which is the restriction of*φ_t_* to 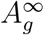. The corollary then follows by applying Theorem 2 to the process 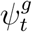.

### Proof of Theorem 3.

As shown in [Edhan et al., 2017], the replicator is an instantiation of Hedge, the exponential version of the multiplicative weights update algorithm. It follows that the replicator attain asymptotic zero regret.

Translating this mathematical result back to the evolutionary setting, this is equivalent to stating that asexual reproduction virtually converges monomorphically to an optimal-in-hindsight genotype *g*\*.

### Proof of Lemma 5.

Focus on a particular allele 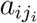 and its attendant 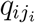. Recall that by Equation (2),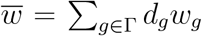, that by Equation (7), 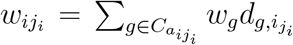, while by Equation (20) the haploid sexual replicator is 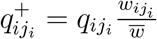.

For 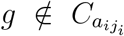, one has 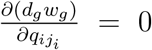. For 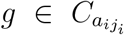, using 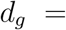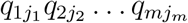 (by linkage equilibrium) yields

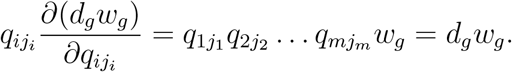

Hence 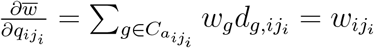. It follows that 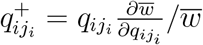.

This is the schema for applying the Baum–Eagon theorem of Equation (24), with *T*_1_ as the transformation and 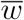 the Lyapunov function.

### Proof of Theorem 4.

By Lemma 5, monotonic mean fitness increase along trajectories is immediate from the Baum–Eagon Theorem.

Any pure strategy profile of the game *W*_Γ_ (corresponding to a point distribution concentrated on a single allele at each locus) constitutes a fixed point of Equation (8), and in fact the only fixed points of this dynamic are pure strategy profiles. However, if *g* is a pure strategy profile of *w* that is not a Nash equilibrium t hen it is not a stable point of the dynamic; using standard dynamics arguments involving nullclines and separatrices, there exists around *g* a neigh-bourhood such that any 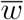-increasing trajectory with initial point in the interior of that neighbourhood eventually leaves that neighborhood.

In other words there is a basin of repulsion around every such point; hence the dynamic cannot converge to non-Nash equilibria points. Convergence will therefore always be to a pure strategy Nash equilibrium point, i.e., a local maximum of the potential 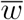.

### Proof of Theorem 5

This is more or less a corollary of Theorem 4. For each pure Nash equilibrium 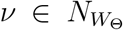, let *B_ν_* be the set of elements in Δ(Γ) that asymptotically converge to 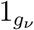. By definition, this forms an asymptotically stable basin of attraction.

### Proof of Proposition 7.

This is an almost direct application of Theorem 3 from [Baum and Sell, 1968]. We can identify *H_ν_* as follows: for any *η* > 0, let *V_η_* be the connected component in Δ(Γ) of 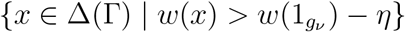 that contains 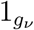.

Let *η*_0_ > 0 be the smallest real number such that 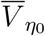, the closure of 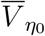, contains another critical point of *w* in addition to 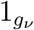. Set 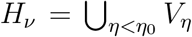. Theorem 3 of [Baum and Sell, 1968] now applies to *H_ν_* to attain the conclusion.

### Proof of Proposition 9.

Suppose that at time *t* the population has mean fitness 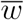. A sexually reproducing sub-population of weight *r* is selected, whose mean fitness is 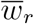, and the complementary asexual sub-population of weight 1 − *r* therefore has mean fitness 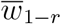, such that 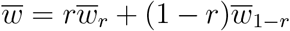. At time *t* + 1, the offspring population of the sexual reproducers has mean fitness 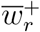, and the offspring population of the asexual reproducers has mean fitness 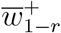.

Since both the *T*_0_ and the *T*_1_ transformations increase mean fitness (except at fixed points), it follows that 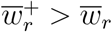 and 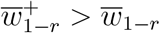. But 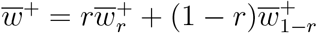, hence 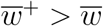.

This argument relies on the *T*_1_ increasing mean fitness monotonically, which rests on Theorem 4, which in turn ultimately relies on Theorem 1. In game theoretic terms, Theorem 1 presumes independent choices of strategies on the parts of the players (as opposed to correlated strategies), hence an initial point of linkage equilibrium neeeds to be assumed. Starting from any such interior point, the trajectory will follow increasing mean fitness until it arrives at a local maximum, which will be a pure Nash equilibrium point.

### Proof of Theorem 6.

Suppose that at time *t* the population has mean fitness 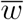. For each partition *λ* ∈ Λ, a sub-population of weight *r_λ_* of reproduction type *λ* is selected, whose mean fitness is 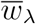 such that 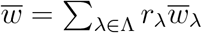. At time *t* + 1, the offspring population of the *λ*- type reproducers has mean fitness 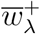, with population mean fitness 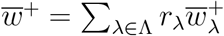

Using similar argumentation as in the proof of Proposition 9, since the *T_λ_* transformations increase mean fitness (except at fixed points) for all partitions *λ*, it follows that 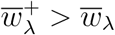 for all partitions *λ*. Hence 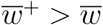.

Starting from any linkage equilibrium point, the trajectory will follow increasing mean fitness until it arrives at a local maximum, which will be a pure Nash equilibrium point.

### Proof of Theorem 7.

In this proof we ask a different question from the usual questions of regret minimisation: instead of asking whether an algorithm attains the same asymptotic rate as the best expert, we suppose that the algorithm converges to the rate of the best expert and ask whether that rate is equal to some exogenous rate.

Let *e*^1^, *e*^2^,… be any environment realisation. Let *i* be a locus. By Equation (8), the alleles in locus *i* are each implementing a replicator equation (with respect to the identical interests game they are playing against the alleles in the other loci). This implies that the reproductive dynamic internal to the locus follows a multiplicative weights updating algorithm with respect to the total environment payoffs, taking into account both external and internal environments. Hence, regret minimisation applies to this dynamic and in the limit the locus attains the average growth rate it would have attained had it implemented the optimal fixed strategy-in-hindsight within Δ(*A_i_*) at all times. In other words, the individual locus attains virtual convergence.

From here the proof proceeds inductively. Suppose that under the true dynamic each locus *i* exhibits the mixed strategy sequence 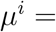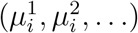 where 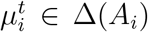 for each *t*. Further, denote by *L* the lim sup average growth rate payoff that is attained under the profile (*μ*^1^, *μ*^2^,…, *μ^m^*) of these strategy sequences (which is equal for each locus).

Let 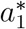 represent the pure strategy of locus 1 whose lim inf attains asymptotically zero regret, as in Equation (14), i.e, that asymptotically does as well as *L*. Locus 2 can then take the perspective of facing an environment consisting of the external environment along with internal environment 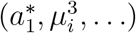, and virtually attain the same payoff with fixed optimal-in-hindsight 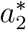, i.e., 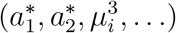 virtu-ally attains *L*.

By induction, the sequence attains 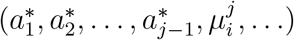. At locus *j*, implement the optimal-in-hindsight 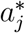 against 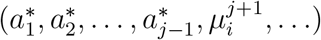 to attain payoff *L* under 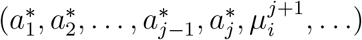.

Continuing by induction, in this way eventually one concludes that population consisting entirely of the genotype 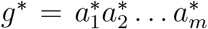 is the optimal-in-hindsight genotype.

### Proof of Proposition 10.

Following the same reasoning as in previous proofs, consider the perspective of locus *i*. The payoff received by locus *i* is equal to what it would gain if all the other loci were to play the pure strategy profile 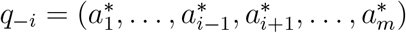.

By assumption the external environment process selecting the realisations 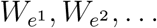 is stationary and ergodic. Since the *q*_−*i*_ pro-file is virtually pure and fixed throughout time, the internal environment 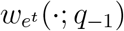 reflects the external environment and is similarly stationary and ergodic. Hence the virtual convergence of locus *i*, which is equivalent to that of an asexually reproducing population under the conditions of a stationary and ergodic environment process, is always to the same payoff. This holds equally true for all loci.

### Proof of Theorem 8.

The proof is the same as the proof of Theorem 5.

### Proof of Proposition 11.

The proof is the same as the proof of Proposition 7.

### Proof of Theorem 9.

By Equation (12), the alleles in the locus are implementing a replicator equation (with respect to the symmetric potential game they are playing amongst themselves). This implies that the reproductive dynamic in the locus follows a multiplicative weights updating algorithm with respect to the total environment payoffs, taking into account both external and internal environments. Hence, regret minimisation applies to this dynamic and in the limit the locus attains the average growth rate it would have attained had it implemented the optimal fixed strategy-in-hindsight within Δ(*A*) at all times.

### Proof of Theorem 10.

Let *e*^1^*, e*^2^,… be any environment realisation. Let *i* be a locus. By Equation (12), the alleles in locus *i* are each implementing a replicator equation (with respect to the symmetric potential game they are playing amongst themselves). This implies that the reproductive dynamic internal to the locus follows a multiplicative weights updating algorithm with respect to the total environment payoffs, taking into account both external and internal environments. Hence, regret minimisation applies to this dynamic and in the limit the locus attains the average growth rate it would have attained had it implemented the optimal fixed strategy-in-hindsight within Δ(*A_i_*) at all times. In other words, the individual locus attains virtual (possibly polymorphic) convergence.

From here the proof proceeds inductively as in the proof of Theorem 7. Suppose that under the true dynamic each locus *i* exhibits the mixed strategy sequence 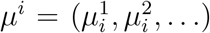, where 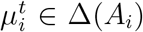 for each *t*. Further, denote by *L* the lim sup average growth rate pay-off that is attained under the profile (*μ*^1^*, μ*^2^,…, *μ^m^*) of these strategy sequences (which is equal for each locus).

Letting 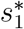 represent the fixed (possibly mixed) strategy of locus 1 whose lim inf attains asymptotically zero regret, as in Equation (14), i.e, that asymptotically does as well as *L*. Locus 2 can then take the perspective of facing an environment consisting of the external environment along with internal environment 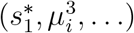, and virtually attain the same payoff with fixed optimal-in-hindsight 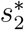, i.e.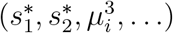, virtually attains *L*.

Continuing argument by induction, in this way eventually one concludes that population consisting entirely of the genotype 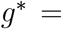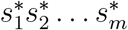 is the optimal-in-hindsight genotype.

## APPENDIX C Proof of Convergence in Potential Games under the Polynomial Multiplicative Weights Update Algorithm

## C.1. Preliminary Setup

Let (*A, u, φ*) be a potential game, where *A* = *A*_1_ … *A_m_* is the set of action profiles, *u* : → ℝ^*m*^ the payoff function, and Φ; : *A* → ℝ the potential. For *x* ∈ *A*, we use *x_i_* to denote the action in *x* of the player *i* and *x*_−*i*_ to denote the actions in *x* of the players apart from player *i*. For simplicity we will assume that |*A_i_*| = *k* uniformly for all players; the extension to the more general case is straightforward. Enumerating the elements of *A_i_*, the *j*-th action of player *i* is 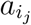.

A mixed strategy of player *i* will be denoted 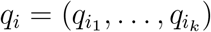, and a profile of strategies *q* = (*q*_1_,…, *q_m_*) ∈ Δ(*A*_1_)×…× Δ(*A_m_*). The ap-plication of a profile *q* yields an expected payoff for player *i* that we will denote *u_i_*(*q*). Given an action profile 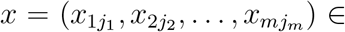 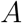 and a profile of strategies *q*, denote

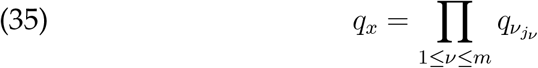

and

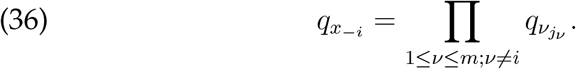

Suppose that each player is applying the multiplicative updates algorithm to update the mixed strategy he uses from one time period to the next. To interpret what is meant by this, we need to specify the payoff player *i* receives for placing weight 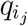 on action 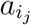 when his overall expected payoff is *u_i_*(*q*).

Suggestively borrowing notation introduced earlier here in the context of alleles, denote 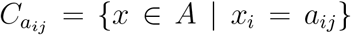, i.e., the set of action profiles with the action of player *i* fixed at 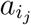. Next suppose that player *i* fixes action 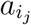 while the other players choose mixed strategies *q*_−*i*_. In this case, denote the expected payoff for player *i* by 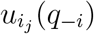, which is

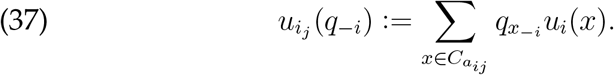

Denote by 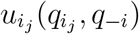 the payoff player *i* receives for placing weight 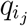 on action 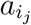 when the other players choose *q*_−*i*_. Using Equation (37), this is

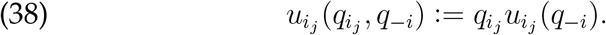

With that we can specify what it means for each player to apply the multiplicative weights updates algorithm for *η* > 0. When *q* is the profile of mixed strategies, player *i* views the tuple 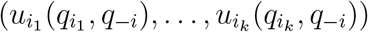. In response, the mixed strategy that player *i* chooses in the next time period is given by

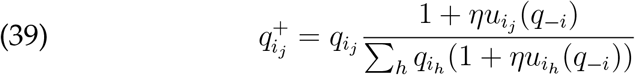

For ease of reading, we will from here express Equation (39) more simply as

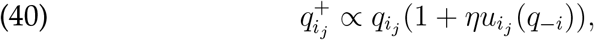

supressing the denominator whose entire purpose is only to ensure that the result is a probability distribution.

### C.2. Proof of Theorem 1.

We prove this first in the special case that the potential game is an identical interests game and that the updating rule is the parameter-free case, i.e., for each *i, j*, *u_i_*(*x*) = *u_j_*(*x*) = Φ(*x*), so that each player gets the same payoff (the potential) for each profile of actions, and Equation (40) becomes

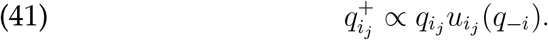

From here most of the work is unravelling of definitions. From Equations (37) and (38) and *u_i_*(*x*) = Φ(*x*) we obtain

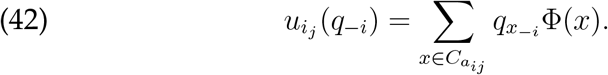

At the same time, the expected payoff of player *i* under *q* is 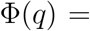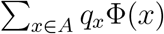. It follows that for each available action 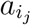,

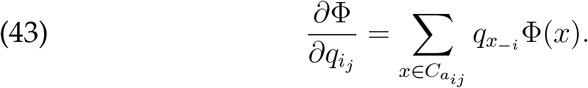

Putting it all together yields

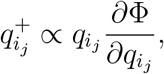

which is exactly what is needed for application of the Baum–Eagon Theorem (since Φ(*q*) is a polynomial function of the various probability weights of *q*). It follows that under this dynamic the value of Φ increases monotonically from one time period to the next, with convergence to a fixed point that is a Nash equilibrium.

Moving on from the parameter-free case, consider next the more general polynomial update rule

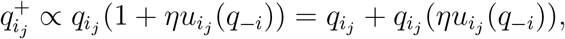

but still maintain the assumption of an identical interest game, i.e., *u_i_*(*x*) = *u_j_*(*x*) = Φ(*x*) for all *i, j*, so that Equation (42) still holds.

Define

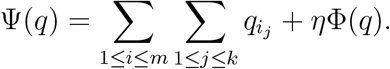

Then

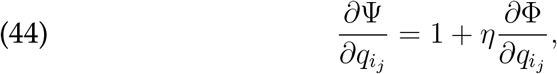

while Equation (43) holds as before, i.e. 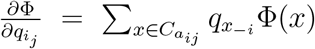.

Putting it all together yields

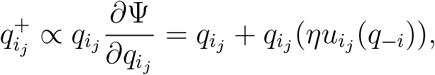

which is sufficient for obtaining the result we seek by appeal to the Baum–Eagon Theorem.

Finally, in greatest generality suppose that the game is fully a potential game as opposed to an identical interests game. In this case, it is well known that the potential game can be decomposed into an identical interests game and a dummy game. That is, *u_i_*(*q*) = Φ(*q*) + *D_i_*(*q*_−*i*_), where Φ is an identical interests game and *D_i_* the payoff to *i* from the dummy game *D*, depends solely on *q*_−*i*_ but does not change at all with changes in *q_i_*.

It follows that 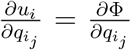: in other words, this case essentially reduces to the case of an identical interests game from the perspective of partial differentiation with respect to the weights of the actions of player *i*. Hence, with minor modifications the same proof as applied earlier applies to the general case; we omit the obvious details.

## APPENDIX D Basins of Attraction and Repulsion

As proved in [Novak and Barton, 2017], using the Baum–Eagon inequality, the haploid sexual replicator dynamic, under fixed fitness conditions, converges to fixed points of the dynamic.^5^

It is clear from inspection of Equation (8) that any pure strategy profile is a fixed point of the dynamic (and conversely the only fixed points are pure strategy profiles); the set of pure Nash equilibria is a proper subset of the set of pure strategy profiles. Details regarding basins of attraction and repulsion, however, need also to be shown.

We will make use of a collection of relative distribution weights (relative to an allele *a_ij_*) 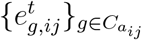. To define this, let 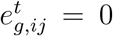 if 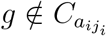. Otherwise, define

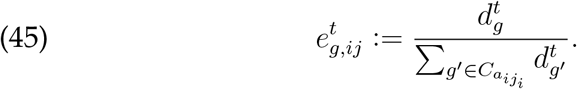

In words, 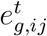 is the weight of gentotype *g* amongst the genotypes containing *a_ij_*. Clearly, 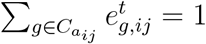.

When linkage equilibrium holds, we can re-express 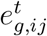 associated with allele 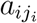 when 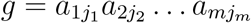 as

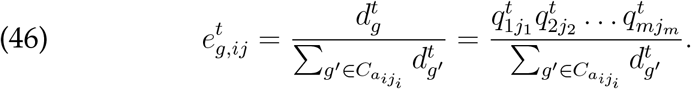

Equation (7), defining the marginal fitness of allele *a_ij_* at time *t* is rewritten in these terms as

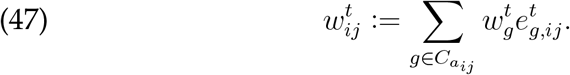

It will be useful to express Equation (47) solely in terms of *q_i_* and *w_g_*. To that end, for a fixed 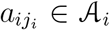, enumerate the elements of 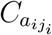 as (*g*_1_,…, *g_l_*). Each such 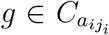 is by definition a string of alleles 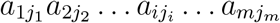 where 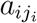 is the same for each 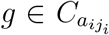 but the other alleles vary from one such genotype to the other.

From Equation (6), under linkage equilibrium, for each such *g* we have 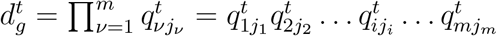. For future reference we will want to use a ‘reduced form’ of this expression, defined as

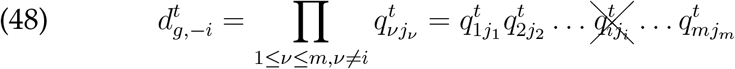

by which we mean the *m* − 1-fold product that does not include 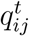 in it. Then we can re-write Equation (46) as

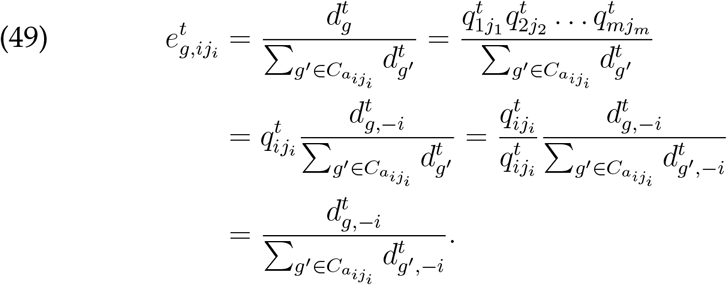

Note that this entirely removes dependence on 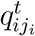; all dependence is on the allelic frequencies of loci *apart* from locus *i*. We can go even further. Denote in vector notation 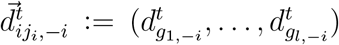, with respect to the eumeration of (*g*_1_,…, *g_l_*) as the elements of 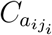. By tracing through the definitions, it becomes clear that the dependence on 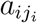 is superfluous. In other words, for 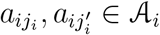, two alleles in locus *i*, one has 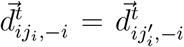. We can therefore denote this vector uniformly as 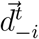

Similarly, 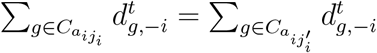 for 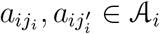. Hence we can choose any one of them and uniformly denote 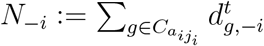.

Continuing with the vector notation, denote 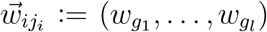 with respect to the enumeration of (*g*_1_, … , *g_l_*) as the elements of 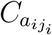. In this we cannot avoid dependence on the specific allele 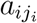.

From here we can rewrite Equation (47) as

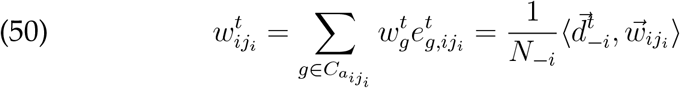

using the vector dot product.

Finally, since the mean fitness 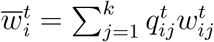 this becomes

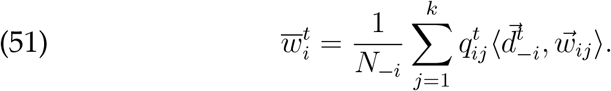

## D.1. Basins of Repulsion Around Non-Equilibrium Points

Suppose that fitness values *w_g_* are fixed over time. Let 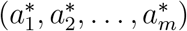 be a pure strategy profile of the game that is not a Nash equilibrium. In the biological interpretation, this corresponds to a population composed solely of genotypes 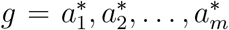, in which case 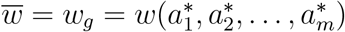.

By definition of a non-equilibrium, there is at least one player/locus 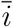 such that, if we write then there is such that if we write 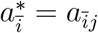 then there is 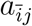 such that 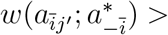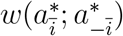. For *ɛ* > 0, construct for each player *i* the mixed strategy 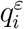 that gives weight 1 − *ɛ* to pure action 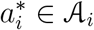 and spreads the rest of the weight *ε* amongst all the other actions in 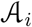.

We claim that for sufficiently small *ε*, the haploid sexual replicator dynamic at the mixed strategy 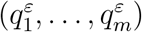 draws away from 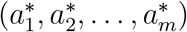

To see this, note that for 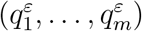 for small *ɛ*, for any *i* and 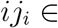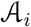, it is the case that 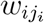 is very close to 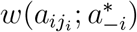. This is because by Equation (50),

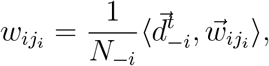

where each element 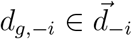, for 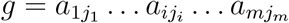 is defined by 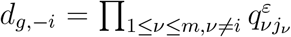. By the definition of 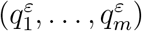, for small *ɛ* the only relevant *d_g,_*_−*i*_ that has appreciable weight is that associated with 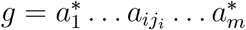, where 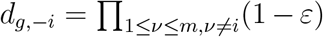. Hence 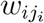 is very nearly 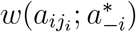.

In particular, 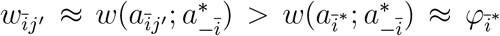. By continuity, for sufficiently small *ε* this strict inequality can be guaranteed to hold, 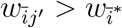.

Since the players are implementing the haploid sexual replicator equation, Equation (8), in updating from one period to the next, 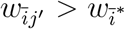 when at time *t* it follows that at time *t* +in the distribution 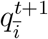 relatively greater weight will be given to 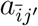 and relatively less to 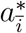. It follows that in the step from *t* to *t* +1 the dynamic moves away from 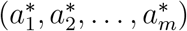.

This is sufficient to conclude that the dynamic must converge to a pure Nash equilibrium.

### D.2. Proof of Proposition 8

Suppose that 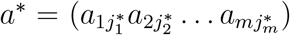 is a pure strategy Nash equilibrium profile of the game.

Let *B* ⊂ *D* be defined as the collection of mixed strategy profiles

(*q*_1_,… , *q_m_*) ∈ *D* such that for all *i*:

1. 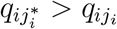 for all 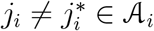 and
2. 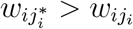 for all 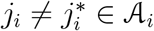.

In words, *B* is the set of mixed strategy profiles such that for each *i*, within the allelic frequency distribution *q_i_* the greatest weight is placed on 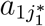 and at the same time the marginal fitness of 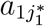 is the highest against all its competing alleles in locus *i*. (Recall that by Equation (7) 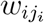 is a function of genotypic density and hence also a function of (*q*_1_,…, *q_m_*).)

We can now proceed to show that *B* is an exponentially stable basin of attraction around *a*\*. To see this, first note that *B* is not empty because the profile *a* * is trivially an element of *B*. By continuity, *B* is then a neighbourhood of *a*\*.

Next, let 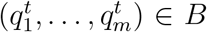 at time *t*. For any *i*, since 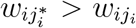 for all 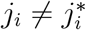, by Equation (8) it follows that at *t* + 1 the weight of allele, 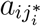 i.e. 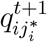 relative to 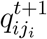 for any other allele, only increases. So the first condition 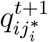 for 1 is satisfied.

Next, since *a*\* is a Nash equilibrium, 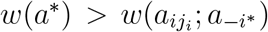 for all 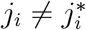. Recall that 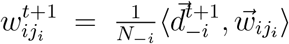. Since the weight 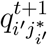 increases relative to 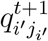 for all *i*^t^, and 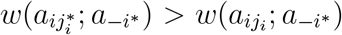 for all 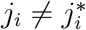, it can only be the case that 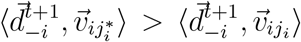, i.e. 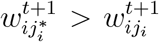. Hence the second condition is satisfied and it follows that 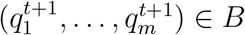.

Finally, since for each *i* the weight 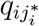 increases relative to any other 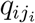 monotonically (and even at an increasing rate), asymptotically starting from any (*q*_1_,…, *q_m_*) ∈ *B* the dynamic converges to *a*\*.

## Footnotes

1 In all models in this paper generations are discrete and non-overlapping, populations are infinite, and no mutation, migration, or genetic drift is included in the models.

2 Strictly speaking we need to consider the lim sup in Equation (14) because the limiting average payoff value might not be well defined.

3 We assume here that there are no position effects.

4 As before, in greater generality it is possible to allow each simplex to be of different dimension and attain the same results. For simplicity of exposition, we restrict here to the special case in which all the simplices are of the same dimension.

5 This result is actually mentioned, without a detailed proof, all the way back in the original paper by Leonard E. Baum and J. A. Eagon [Baum and Eagon, 1967].

